# Fast prediction of acidic amino acid sidechain conformations for cryo-EM modeling

**DOI:** 10.64898/2026.07.12.738023

**Authors:** Georgios Kolypetris, Amina Djurabekova, Jonathan Lasham, Luka Simsive, Janet Vonck, Vivek Sharma

## Abstract

Cryogenic-electron microscopy (cryo-EM) has revolutionized the field of protein structural biology. The structures of large membrane proteins are now routinely determined by cryo-EM to near atomic resolution. However, in the medium resolution range of cryo-EM maps (>∼2 Å), negatively charged sidechains of acidic residues are not well-resolved due to the negative electrostatic potential of the region. This may lead to incorrect sidechain models for residues like glutamic acid or aspartic acid that are central for proton transfer activity in various respiratory and photosynthetic enzymes. We previously proposed that the acidic residues with weak or non-existent cryo-EM density can be modeled to represent their low proton affinity conformations. Here, we tested this hypothesis on a larger data set of acidic amino acid residues in two high-resolution respiratory complex I structures. By using faster sidechain modeling and proton affinity prediction tools, we created a workflow that generates sidechain conformations of selected amino acid residues. We validated the sidechain conformation predictions by Q-score analysis and atomistic molecular dynamics simulations in different charged states. The proposed workflow provides a way to rapidly obtain sidechain conformations of acidic residues with weak cryo-EM densities and can be integrated into the existing cryo-EM modeling pipelines to speed up sidechain rotamer prediction.

## Introduction

The photosynthetic and respiratory enzymes catalyze proton transfer reactions as part of their energy generating functions [1–3]. The high-resolution 3D structures from X-ray crystallography and more recently by cryogenic-electron microscopy (cryo-EM) have provided detailed atomic-level insights of the protein interior, such as the hydrophilic regions with bound water molecules [4–6] (Fig. 1). These hydrated polar domains are thought to catalyze proton transfer reactions, often linked to some photo or redox activity and/or conformational transitions [1, 7]. The diffusion of protons occurs along a chain of water molecules and sidechains of polar amino acid residues via a Grotthuss-like proton transfer mechanism [8]. The conformational mobility of proteins and associated proton transfer dynamics can be studied with the sophisticated computer simulation techniques, and also in a quantitative framework by calculating reaction energies, including driving forces and activation energy barriers [9–13]. Conventional classical molecular dynamics (MD) simulation approaches are able to capture the nanoseconds to microseconds dynamics of proteins and water molecules at atomic resolution [14, 15], but proton transfer reactions are not explicitly modeled. MD simulations performed in different charged states, determined by empirical p*K*_a_ calculations [16] or continuum electrostatics [17, 18], revealed functional dynamics of amino acid sidechains [19, 20] and hydration/dehydration effects [14, 19, 21–23]. Recent improvements in speed and accuracy of discrete and continuous constant pH classical MD simulations have allowed the exploration of protonation dynamics at extended timescales [24–26]. On the other hand, quantum chemical approaches, such as the DFT-based QM/MM calculations [27, 28], can be utilized to simulate explicit proton transfer reactions at short time and length scales.

**Figure 1.**
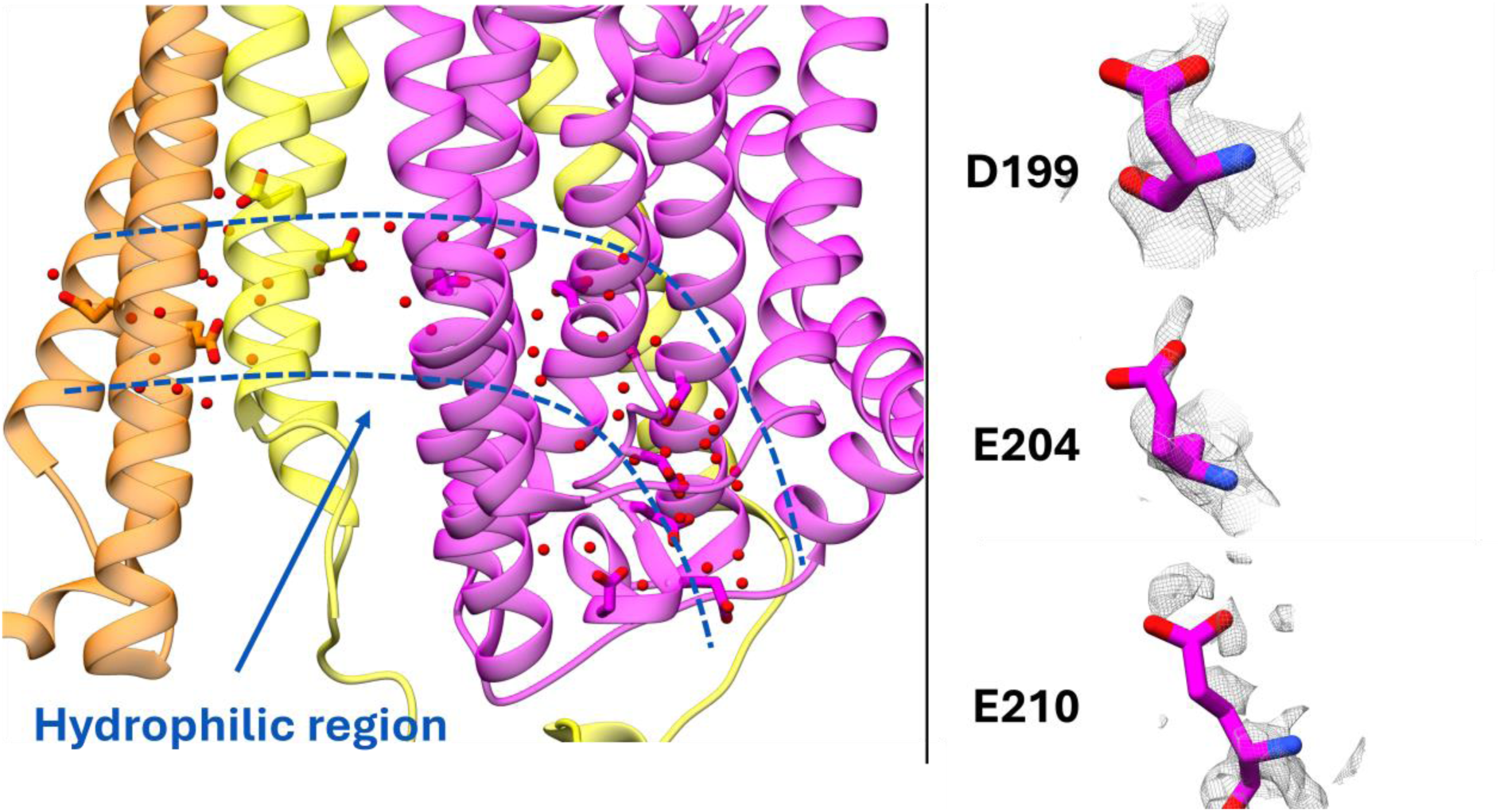
Acidic amino acid sidechains and water molecules in hydrophilic domains of proteins. (Left) Hydrated water-filled polar domain (blue dotted lines) called the E channel of proton transfer in mouse respiratory complex I structure (PDB 8OM1). (Right) Cryo-EM density (grey mesh) and atomic model based on it (sticks) shown for selected acidic amino acid residues from PDB (EMDB) 8OM1 (16965) and 7O71 (12742). The selected examples of a good (D199), a weak (E204) and a nearly non-existent (E210) cryo-EM density map are shown.

Crucial to all the computational modeling and analysis described above is the high resolution of the protein structure data, on which reliable simulation models are constructed. The accuracy of the output from computer simulations depends to a good extent on the quality of the 3D structure data used. MD simulations performed on low resolution data produce noisier trajectories that display higher RMSDs of the protein (see e.g. [29, 30]). If the atomic model obtained from X-ray crystallography or cryo-EM is of such a low resolution that the backbone or sidechains are entirely missing, computer simulation approaches cannot always identify the native protein conformations, due to limited sampling times and force-field accuracy. It has been observed that classical MD simulations performed on homology models (especially based on low sequence identity) often lead to model deterioration, instead of stabilizing to the native state [31] (see also [32]). Although there is evidence in which the computer modeling of low-resolution structural data yielded functionally relevant conformations [33–35]. Nevertheless, it is recommended to build and simulate protein models on high-quality structure data [36].

Cryo-EM has revolutionized the field of protein structure determination [37, 38]. Despite its remarkable success, high-resolution cryo-EM maps display regions where the density is either weak or completely non-existent, which can lead to suboptimal modeling of amino acid sidechains, especially acidic residues (see below). There are several known reasons for the cryo-EM density to be weaker [39–41]. If the region is conformationally mobile (e.g. a loop), the image averaging procedure does not yield a sufficiently sharp density map required to construct a reliable atomic model. Second, prolonged electron exposure leads to higher levels of inelastic scattering and chemical perturbations, which can result in loss of density (also called radiation damage). A third phenomenon that occurs in low-to-medium resolution cryo-EM maps (>∼2 Å), is that the scattering amplitude of anionic regions dives to negative values, resulting in weak or completely non-existent density (Fig. 1; e.g. [41, 42] and references therein), thereby preventing accurate atomic modeling from the cryo-EM data.

Building upon the last notion, we have recently proposed that the sidechain conformation of an acidic residue with weak or missing cryo-EM density can be modeled to represent its low proton affinity states (e.g. pKa < 7 at pH 7)[19]. We applied this strategy to remodel the sidechain conformations of selected acidic amino acid residues whose densities were found to be either weak or non-existent [19]. We found that remodeling the sidechain into a low proton affinity conformation also improved the proton affinity predictions for those amino acid residues that were resolved well in the cryo-EM data [19].

In this work, we applied the strategy to a larger data set of amino acid residues from two protein complexes, and automated the process of conformation generation and selection. The developed workflow results in an ensemble of suitable sidechain conformers for acidic amino acid residues. We validate our proof-of-concept in several different cases where acidic residues are found to be either protonated or deprotonated based on cryo-EM density and structural analysis. The results from the workflow are also complemented by Q-score analysis [43] and atomistic MD simulations, which yield alternative sidechain conformations that overlap with the predictions from the modeling. We envisage that this fast and simple protocol could be useful for cryo-EM modeling and refinement pipelines.

## Results

### Workflow

The workflow (described in Fig. 2) employs existing methods and tools, and the result is a desired pipeline that provides an ensemble of sidechain conformations of selected acidic amino acid residues (aspartic and glutamic acids) for which the cryo-EM density is either weak or entirely missing. The cryo-EM densities of basic amino acid residues such as lysine or arginine, which are also functionally important in proteins catalyzing proton transfer reactions [23, 44–46], are usually well-resolved and therefore are omitted from our analysis. The workflow employs MODELLER [47] software to generate a user-defined number of sidechain conformations for target residues (default 100 conformers). During conformer generation, coordinates of the protein remain fixed and only the dihedral (*d* = C-Cα-Cβ-Cγ) of the selected acidic residue is varied. A fast pKa calculation is performed on each conformer using empirical PROPKA software [16]. Since the acidic amino acid residues with weak cryo-EM density are likely to be in their anionic charge states (i.e. deprotonated), the conformers are selected based on a user defined pKa cutoff (default ≤ 7). Additionally, the workflow provides a way to identify a representative sidechain conformation from the ensemble using user-defined clustering approaches.

**Figure 2.**
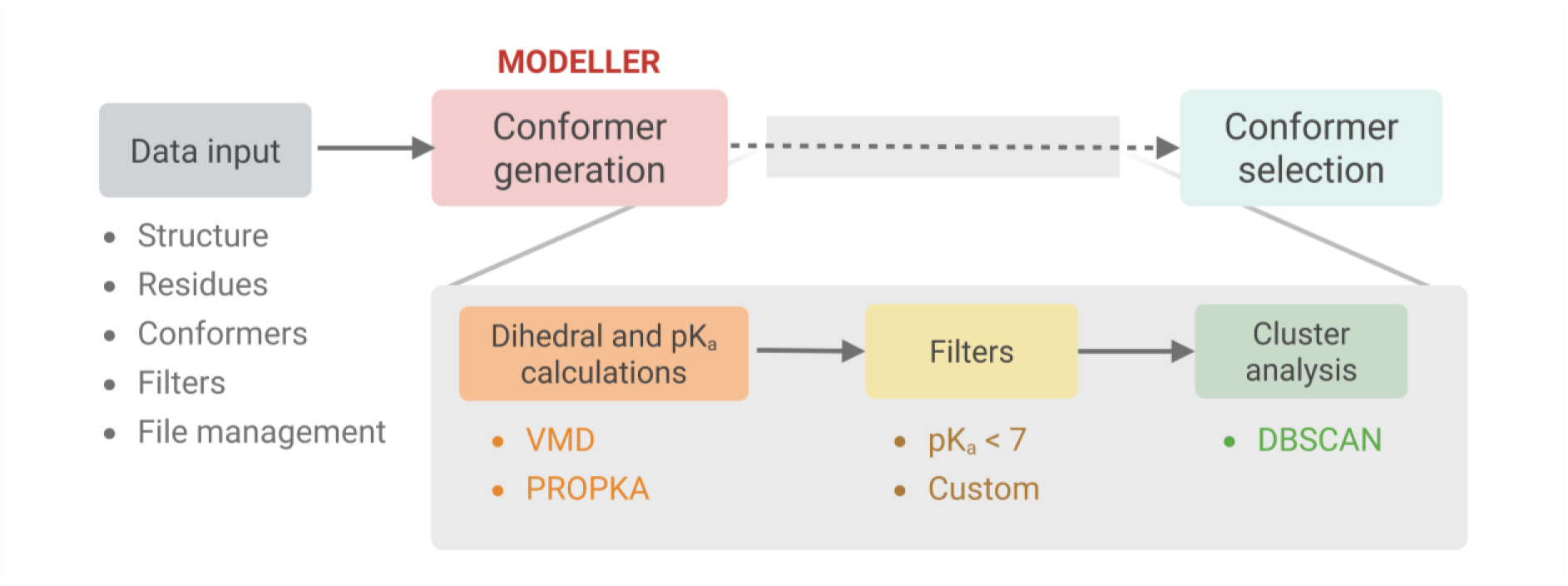
Computational pipeline. Workflow for sidechain conformation generation, pKa calculation and selection of low proton affinity sidechain conformers.

### Testing of the workflow

In order to probe the functionality and accuracy of the workflow, we focused on respiratory complex I, which uniquely consists of several functionally important glutamates and aspartates in the middle of its ∼150 Å long membrane domain [48]. These acidic residues are responsible for long-range proton transfer in complex I. We analyzed several acidic residues from the two high-resolution cryo-EM structures of mitochondrial complex I, from the yeast *Yarrowia lipolytica* (*Yl*, PDB 7O71, ∼2.4 Å, best local resolution ∼2.1 Å, [14]) and from *Mus musculus* (*Mm*, PDB 8OM1, 2.4 Å, [49], see also Table 1, and Methods). See Table 1 for a list of all amino acid residues studied in this work.

**Table 1.** Amino acid residues studied. *Yl* and *Mm* denote *Yarrowia lipolytica* and *Mus musculus*, respectively. Homologous residues from complex I of the two organisms are listed in a single line. EMDB-derived average residue Q-scores are shown. Values in parentheses are the Q-scores derived from ChimeraX by aligning the model (PDB structure) to the map.

| Protonation state based on cryo-EM density | *Yl* PDB 7O71 |  | *Mm* PDB 8OM1 |  |
| --- | --- | --- | --- | --- |
|  | chain/residue | Q-score | chain/residue | Q-score |
| Protonated | 1/E196 | 0.727 (0.839) | H/E192 | 0.729 (0.867) |
|  | 3/D67 | 0.779 (0.873) | A/D66 | 0.724 (0.800) |
|  | 1/E147 | 0.750 (0.827) | H/E143 | 0.709 (0.827) |
| Deprotonated | 1/D203 | 0.659 (0.764) | H/D199 | 0.739 (0.882) |
|  | 1/E208 | 0.477 (0.494) | - | - |
|  | - | - | H/E206 | 0.678 (0.740) |
|  | - | - | H/E204 | 0.583 (0.614) |
|  | 1/E210 | 0.343 (0.198) | - | - |
|  | 1/E206 | 0.644 (0.732) | H/E202 | 0.638 (0.710) |
|  | 1/E231 | 0.656 (0.680) | H/E227 | 0.560 (0.511) |
|  | 3/E69 | 0.694 (0.735) | A/E68 | 0.692 (0.755) |
|  | L/E30 | 0.665 (0.32) | K/E34 | 0.696 (0.798) |
|  | L/E66 | 0.650 (0.646) | K/E70 | 0.708 (0.781) |
|  | 5/D397 | 0.502 (0.652) | L/D393 | 0.460 (0.397) |

### Modeling the sidechain conformations of acidic residues with a high-quality cryo-EM map

We first considered acidic amino acid residues that are likely to be in their anionic charge states, stabilized in part by the positively charged lysine and/or arginine residues. The sidechains are often well-resolved in such cases. For example, D199 in *Mm* complex I and homologous D203 in *Yl* complex I show reasonably high Q-scores (Table 1), and have well-defined cryo-EM densities, which lead to their unambiguous atomic models (see Fig. 3). Positively charged arginine residues (R279 and R302, Fig. 3) are present near to these acidic sidechains; the electrostatic interactions between the residues lead to their sidechain stabilization, as well as charge neutralization, which yields sharper cryo-EM densities, thereby allowing reliable atomic model construction. We find that our Modeler-Propka workflow correctly predicts the sidechain conformations of these acidic residues, closer to the conformations observed in high-resolution cryo-EM structures (Fig. 3). In *Mm* complex I models, D199 shows two sidechain clusters: a smaller group with dihedral *d* ranging from -100° to -50° and a larger cluster with *d* ∼ +50° to +100°. The low proton-affinity small-sized cluster captures the structural conformation of D199, whose sidechain is stabilized by the positively charged arginine R279 (Fig. 3). In the other cluster, the sidechain of D199 points away from the arginine residue and displays an upshifted pKa.

**Figure 3.**
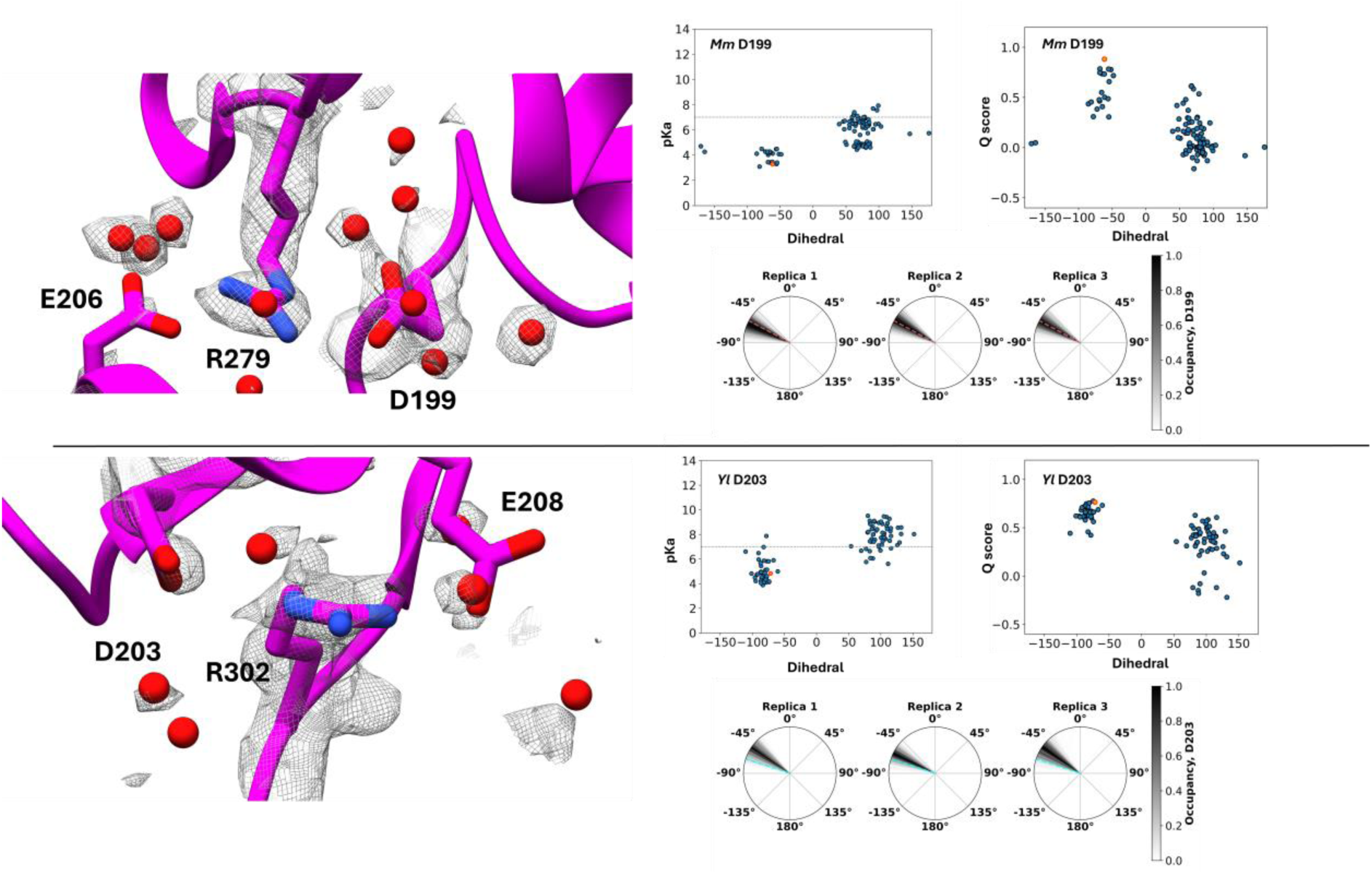
Modeling anionic sidechains with well-defined cryo-EM densities. The cryo-EM map (grey mesh) and sidechain models based on the density are displayed in left panels (top and bottom – mouse and yeast complex I, respectively). Right panels display pKa vs dihedral angle (C-Cα-Cβ-Cγ) and Q-score vs dihedral angle plots for homologous acidic residues (D199 and D203 from mouse and yeast complex I, respectively). The occupancy of the dihedral angle from MD simulations are shown in radial heatmaps with three independent replicas. pKa and Q score of structural conformations are marked with orange dots. The dotted line in pka/dihedral plot corresponds to the pKa value of 7. The dihedral angle corresponding to structural conformations are displayed with pink and cyan dotted lines for *Mm* and *Yl* complex I, respectively. *Mm* – *Mus musculus* (mouse) and *Yl* – *Yarrowia lipolytica* (yeast) mitochondrial complex I.

The acidic residue D203 in *Yl* complex I (homologous to D199 of *Mm* complex I) also displays two similar dihedral clusters (Fig. 3). In the cluster encompassing the structural conformation, the sidechain of D203 points towards R302, whereas when rotated to a dihedral angle in the range *d* ∼ +50° to +100°, it points away from the arginine residue resulting in an upshifted pKa value (Fig. 3). Interestingly, this alternative high-pKa ensemble resembles the sidechain arrangement observed in the cryo-EM structure of *Yl* complex I obtained under *turnover* conditions [50], suggesting that this alternative arrangement, identified by our fast and simple modeling procedure, may have mechanistic relevance.

### Estimating the accuracy of the modeled sidechains

Among other metrics [51, 52], Q-score is one that describes the resolvability of atoms in a cryo-EM map [43]. A higher value of Q score (closer to 1) means a better fit of the sidechain atomistic model to the experimental density. We list the Q-scores obtained directly from the PDB and we also calculated the same for the modeled sidechains against the experimental cryo-EM maps using ChimeraX [53] (EMD-12742 and EMD-16965, see Table 1). We find that in the case of D199 in *Mm* complex I (and homologous D203 in *Yl* complex I), for which the quality of the cryo-EM maps and the resulting structural model are very high, the reported and calculated Q-scores are also excellent (see Table 1). For the modeled sidechain conformations belonging to the dihedral angle space -50° to -100°, the Q-scores obtained are relatively higher than the ones in alternative conformations (+50° to +100°, Fig. 3) suggesting that the former dihedral angle range correctly represents the sidechain position, also commensurate with the electrostatic stabilization discussed above.

To further probe the accuracy of the modeled sidechains, we analyzed the classical MD simulation data of *Yl* and *Mm* complex I [14, 54, 55] and find that the sidechains of anionic D199 and D203 remain stable in microsecond timescales (Fig. 3). Importantly, the dihedral values obtained from simulations fall into the same range as those observed with our Modeler-based modeling protocol. The data from simulations consolidate our modeling protocol that can reproduce the structurally observed conformations and also generate additional conformers representing the cryo-EM density.

### Modeling protonated acidic amino acid residues with good cryo-EM density

We next analyzed the acidic residues E143 and E192 in *Mm* complex I, and homologous E147 and E196 in *Yl* complex I. Since the cryo-EM density is well resolved in these cases (Fig. 4, see also Table 1 for Q-scores) and because there are no charge-stabilizing residues (or ions) in their vicinity, these acidic amino acid residues could be considered partly protonated (charge neutral), in contrast to the anionic cases discussed above (D199 and D203). As shown in Fig. 4, our workflow leads to several sidechain conformers (mostly in the 0° to 180° range) that also cluster around the conformations observed in the cryo-EM structures. The modeled conformers consistently show pKa values ≥ 7, as opposed to the case of D203 and D199 (Fig. 3). Even though the Q-scores of the sidechains derived by mapping PDB structure onto the cryo-EM map are often higher, the modeled sidechains do show comparable Q-scores; for example, at *d* ∼ -50° for E192 and at *d* ∼ +150° for E143 in *Mm* complex I (see Fig. 4). Analysis of dihedral angles obtained from MD simulations also corroborates the modeled sidechain conformations, with alternative sidechain arrangements similarly populated (SI Fig. 1).

**Figure 4.**
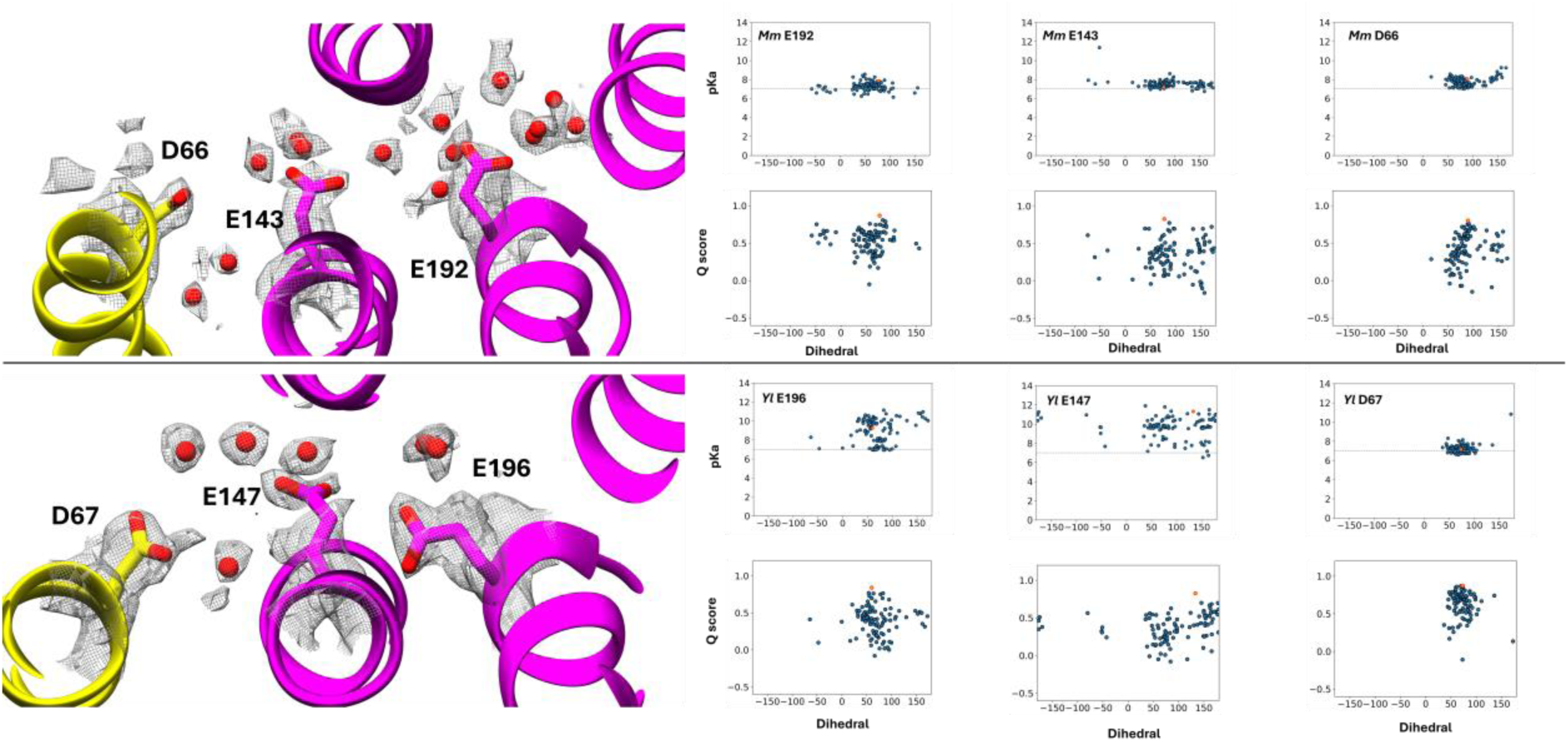
Modeling sidechains of protonated acidic residues with well-defined cryo-EM densities. The cryo-EM map (grey mesh) and sidechain models based on the density are displayed in left panels (top and bottom – mouse and yeast complex I, respectively). Right panels display pKa vs dihedral angle (C-Cα-Cβ-Cγ) and Q-score vs dihedral angle plots for selected acidic from mouse and yeast complex I. pKa and Q score of structural conformations are shown as orange dots, whereas dotted lines in pKa/dihedral plots correspond to pKa of 7. *Mm* – *Mus musculus* (mouse) and *Yl* – *Yarrowia lipolytica* (yeast) mitochondrial complex I.

The cryo-EM densities of homologous D66 and D67 (*Mm* and *Yl* complex I respectively) are also relatively well-resolved (see Table 1), suggesting that they too are charge neutral. The Modeler/Propka protocol is able to pick the sidechain conformations with pKas ≥ 7, in good agreement with the density-based assignments (Fig. 4). The Q-score analysis and sidechain dynamics observed in classical MD simulations also support the predicted conformations (Fig. 4 and SI Fig. 1).

So far, our modeling protocol shows that the sidechain conformers from Modeler when combined with the faster pKa prediction tool Propka, yields conformations that resemble the cryo-EM models and density-based charge-state predictions. The predicted conformational states show reasonable Q-scores and are also corroborated by MD simulations. We note that while structural conformations are well-predicted, alternative sidechain arrangements are also populated. Currently, there are no higher resolution cryo-EM maps available to confirm the alternative positions found by our methodology, but in the interim the predicted conformational ensemble may turn out to be useful for cryo-EM refinement procedures. Also, since the conformation generation and pKa calculation via the Modeler/Propka protocol are rapid (minutes), this fast approach can be integrated into modeling packages (e.g. ChimeraX and Coot [53, 56]) to obtain inexpensive conformational ensembles of acidic residues with challenging densities.

### Acidic sidechains with weak or missing cryo-EM densities

Now that our workflow can produce reliable sidechain conformations of residues with well-resolved cryo-EM densities, we next tested it on the challenging cases where cryo-EM density of the sidechain is weak or entirely missing. The loss of density reduces the reliability of the atomic model, a limitation that current refinement procedures in cryo-EM modeling pipelines cannot fully address. Leveraging on the fact that the residue is likely deprotonated (that’s why density is missing!), we analyzed the pKa *vs.* dihedral relationships of several such acidic residues from the two high-resolution cryo-EM structures.

We start with the acidic residues E202, E204 and E206 of *Mm* complex I (Figs. 3 and 5) and corresponding E206, E208 and E210 from *Yl* complex I (Figs. 3 and 6). These residues reside on a conserved and functionally important loop in respiratory complex I [57, 58]. In the *Mm* complex I structure, E206 is modeled closer to the ion-pair R279/D199 (Fig. 3). In contrast to the compact dihedral clusters observed for anionic D199, a somewhat scattered distribution of sidechain dihedrals is observed for E206 (Fig. 5). The structural conformation is found among the modeled data points, which all display pKa values well below 7, suggesting that anionic E206 can be stabilized by the positively charged arginine residues in its vicinity (R279 and also R238 and R134 (not shown) in *Mm* complex I, Fig. 3). Interestingly, several sidechain conformations with dihedral angle -150° to -100° display Q-scores comparable to the Q-score of the structural model. The highly-conserved acidic residue E208 from *Yl* complex I resides next to a conserved arginine (R274, Fig. 3) suggesting a possible electrostatic stabilization of the two charged sidechains. However, cryo-EM density of E208 is rather weak (Fig. 3) commensurate with the low value of reported and calculated Q-scores (Table 1, see more below). With our modeling protocol, we can reproduce the structural conformation of E208, however, several other low pKa conformational clusters with comparable Q-scores are also obtained (Fig. 6). In both cases E206 (*Mm* complex I) and E208 (*Yl* complex I), for which several low-pKa (anionic state) conformations are found, we obtained Q-scores comparable to the ones from experimental model, suggesting that the predicted sidechain arrangements are valid solutions for the given cryo-EM density.

**Figure 5.**
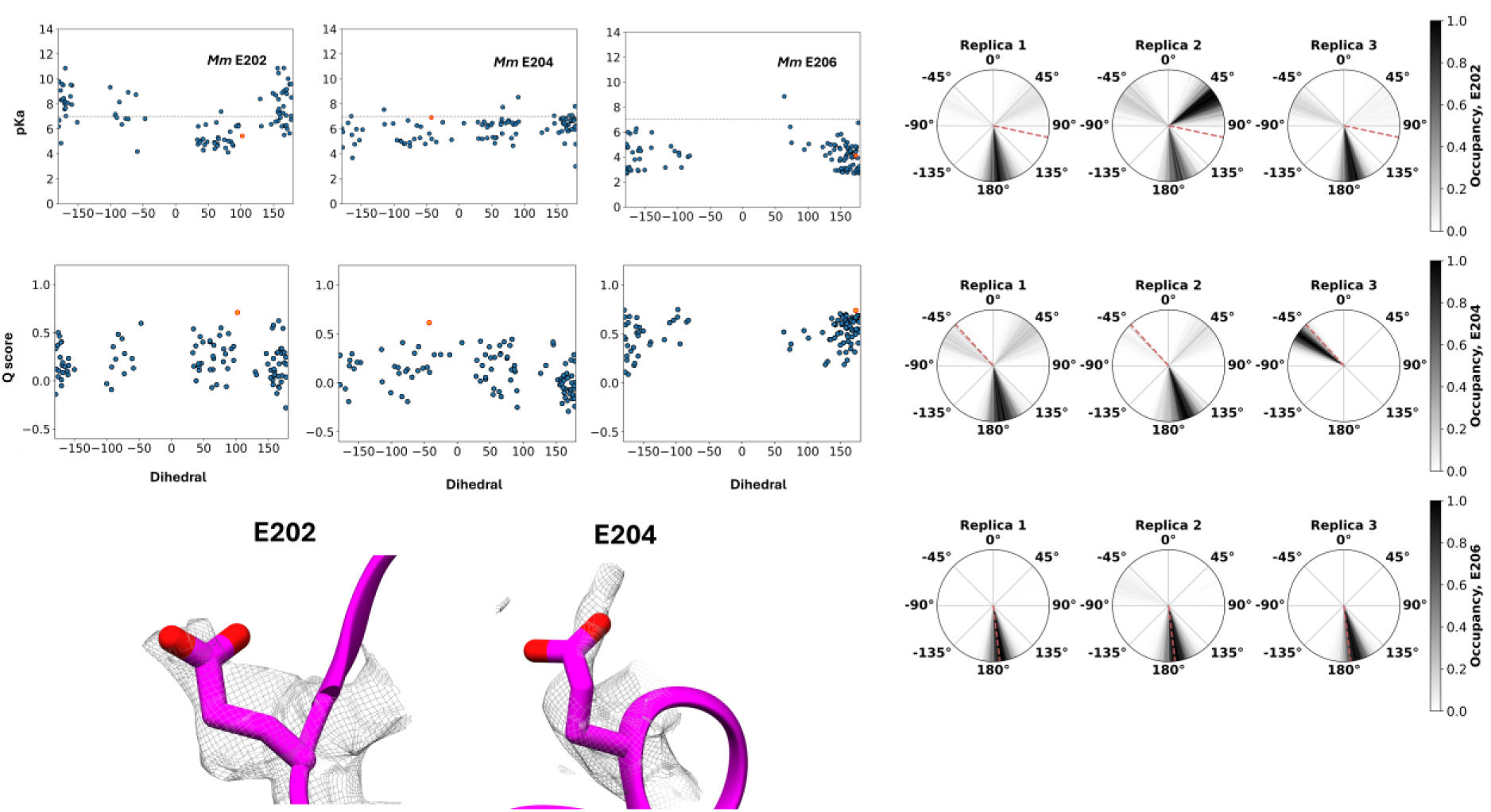
Modeling sidechains of anionic acidic residues in mouse (*Mm* – *Mus musculus*) complex I with weak cryo-EM densities. The left panels display pKa vs dihedral angle (C-Cα-Cβ-Cγ) and Q-score vs dihedral angle plots, and cryo-EM density/atomic model for selected amino acid residues. The right panels show MD simulation based dihedral angle occupancy as radial heat maps for three different simulation replicas (structural conformation is represented by dotted pink line). pKa and Q score of structural conformations are shown as orange dots in plots, whereas grey dotted line corresponds to pKa value of 7.

**Figure 6.**
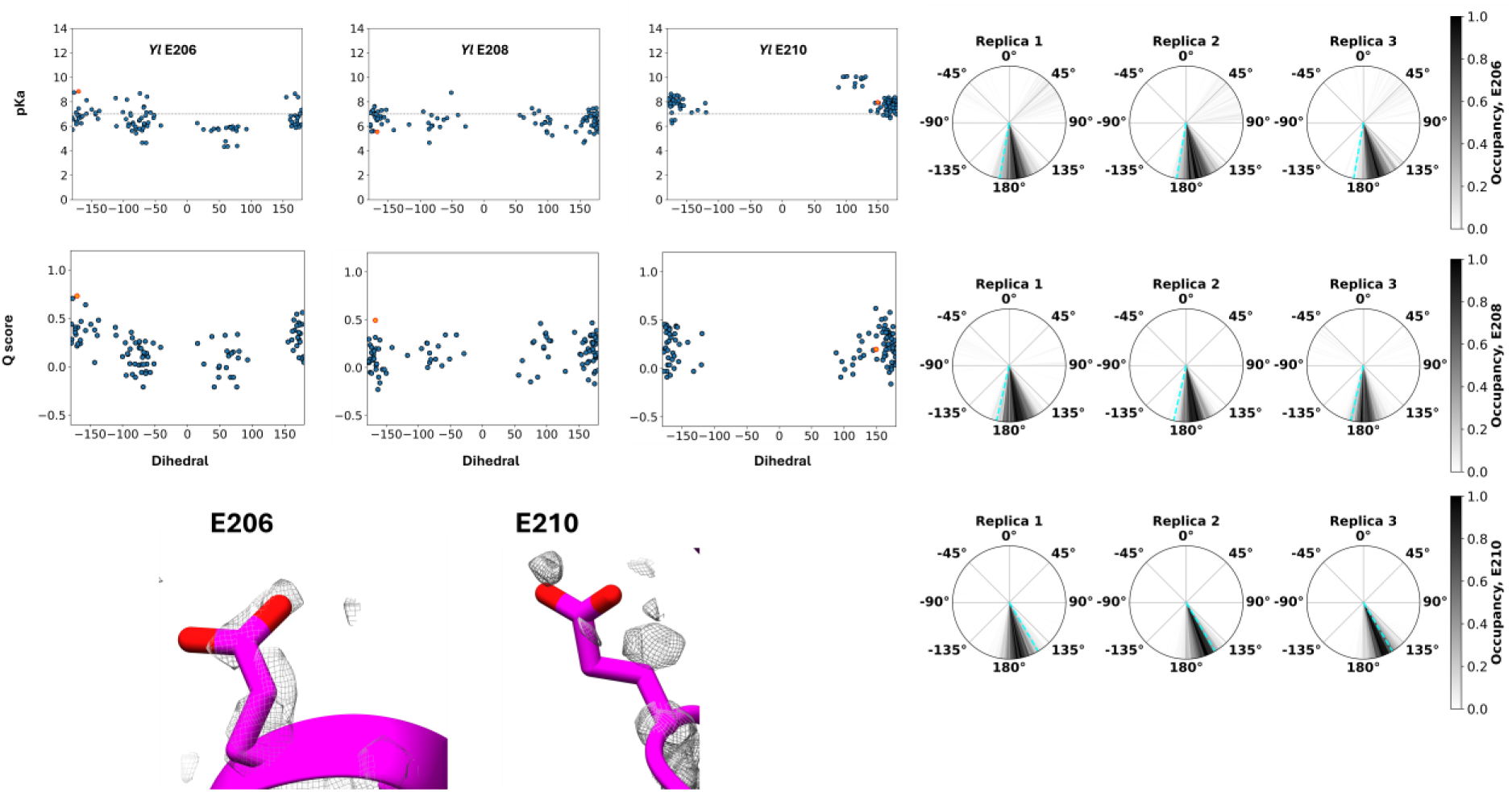
Modeling sidechains of anionic acidic residues in yeast (*Yl* – *Yarrowia lipolytica*) complex I with weak cryo-EM densities. The left panels display pKa vs dihedral angle (C-Cα-Cβ-Cγ) and Q-score vs dihedral angle plots, and cryo-EM density/atomic model for selected amino acid residues. The right panels show MD simulation based dihedral angle occupancy as radial heatmaps for three independent replicas (structural conformation is represented by dotted cyan line). pKa and Q score of structural conformations are shown as orange dots, whereas grey dotted line corresponds to pKa value of 7.

Both E202 and E204 display several conformational clusters with low pKa and different from the structurally modeled conformations (see Fig. 5). Similarly, for E206 in *Yl* complex I, several low pKa conformations are obtained (Fig. 6). The Q-score analyses on modeled sidechains show that the models display Q-scores comparable to the structural values (Fig. 5 and 6, Table 1). We do note however that in some cases the Q-score of the structural conformation is still the highest, whereas the predicted conformations display relatively lower Q-scores. The reason for this discrepancy could be overfitting (fitting to the density noise). Interestingly, for both E202 and E204, MD simulations are able to identify sidechain conformations that resemble the sidechain arrangements seen in models. Similarly, for *Yl* complex I E206, the low pKa conformers in dihedral angle range -100° - +100° are also observed in atomistic MD simulations, albeit weakly. The acidic residue E210 from *Yl* complex I shows a stable conformation resembling the structurally predicted sidechain arrangement, in part due to its electrostatic interaction to a conserved arginine residue. A similar analysis was performed on several other acidic residues present in the membrane domain of *Mm* and *Yl* complex I (see SI Figs 2 and 3). The combined data shows that our workflow can predict several low proton affinity conformations in addition to the structurally observed poses. These alternative conformations that represent the anionic nature of the sidechain can be considered possible solutions to the weak (or non-existent) cryo-EM density map.

## Discussion

The last decade has seen a massive surge in high-resolution protein structures. Membrane protein structures, in particular, which have been challenging to characterize with conventional X-ray crystallography, are now routinely studied with cryo-EM. This has become possible due to dramatic developments at both the hardware and algorithmic levels in cryo-EM-based protein structure determination [37]. Among several areas that are witnessing rapid developments, the accurate modeling of amino acid sidechains from the cryo-EM density data is one challenge that can benefit from computational modeling [19]; in particular the sidechains of acidic amino acid residues, which are hard to observe in the cryo-EM maps of medium resolution [42, 59]. The enhancements in high-performance computing hardware and simulation software [60, 61], which now allow for large, long-timescale simulations, together with constantly improving force-field accuracy [62], can help identify sidechain conformations of acidic amino acid residues [19]. However, atomistic MD simulations are still slow to use in cases where the protein of interest is large; for instance, a full-sized atomistic model of ∼1 MDa respiratory complex I in membrane-water environment comprises a whopping ∼1.3 million atoms, which poses a significant computational challenge for simulation sampling. Therefore, fast and approximate methods are needed that can identify functionally relevant conformations of amino acid sidechains in weak density regions. We here present a workflow that combines two readily available and well-established tools, Modeler software [47] to generate sidechain conformers of acidic amino acid sidechains, and the empirical pKa calculation tool Propka [16] to filter out conformations that display user defined pKa criteria.

By applying the workflow on cases where the sidechain positions are known with high certainty (sharp cryo-EM density), we validate the applicability of the Modeler/Propka combination in predicting the sidechain conformers. We next considered those amino acids for which the atomic model is unreliable due to weaker cryo-EM density. We find that several unique low pKa conformers can be predicted by the Modeler/Propka pipeline, distinct from the conformations obtained based on cryo-EM. Due to the lack of higher resolution structural data (where even anionic sidechains would be visible [59, 63], unless chemically truncated or conformationally mobile), the accuracy of predicted conformations is hard to validate. Nevertheless, Q-score analysis and classical MD simulations performed in different charge states support the Modeler/Propka-based conformer predictions. We note that for completely missing cryo-EM densities of acidic residues, the Q-score is not a useful metric in the current context. Nevertheless, for weak cryo-EM density situations, Q-scores of sidechains modeled in alternate conformations are found to be comparable to the Q-scores of structural models.

In the current implementation, sidechain modeling is achieved with Modeler software, but alternative approaches such as Dunbrack’s conformer libraries [64] can be utilized to generate sidechain conformations. The Propka pKa calculation method is fast and reliable, especially for buried residues and has consistently ranked high in several pKa prediction tests [65, 66]. More thorough pKa prediction approaches such as free energy perturbation [67], constant pH [24, 26] or QM-based methods [68] can be utilized to accurately derive proton affinities, but the computational burden of these methods is high. An additional possibility is to use ML-based pKa prediction tools [69, 70] and apply those on Modeler-derived sidechain conformational set.

## Methods

The sidechain conformers were generated and analyzed using a computational pipeline as shown in Fig. 2 (see also [71]). The pdb file of the cryo-EM structure was used as input and sidechain conformers of selected amino acid residues (acidic residues) were generated using Modeler [47] software. Each of the output files (in pdb format) consisted of a single unique sidechain conformation of the selected acidic amino acid residue, whereas the rest of the protein was maintained in its structural conformation. PropKa [16] method was used to calculate the pKa of the selected amino acid residue, whereas the corresponding dihedral angle C-C_α_-C_β_-C_γ_ (common to both Asp and Glu residues) was calculated using VMD [72]. Conformers not yielding pK_a_ values are not taken into consideration. The scripts related to entire workflow are available at https://github.com/vsharma-cbg/TARCS. The MD simulations of mouse and yeast complex I structures are described in [14, 55]. The simulations of protein complexes [14, 54, 55] were performed in membrane-solvent environment using CHARMM force field [73–75] and Gromacs [76] simulation engine.

## Author contributions

GK performed research, analyzed data, prepared figures

JL performed research, analyzed data

AD analyzed data, prepared figures

LS performed research, analyzed data, prepared figures

JV analyzed data

VS designed research, performed research, analyzed data and wrote the paper with input from all co-authors.

## Acknowledgements

VS acknowledges research funding from the Finnish Society of Sciences and Letters, the Research Council of Finland, the Jane and Aatos Erkko Foundation, the Sigrid Jusélius Foundation, and the University of Helsinki. We are thankful to the Center for Scientific Computing (CSC) Finland for high-performance computing resources. LS is supported by SciDoc-CHEMS doctoral programme of the University of Helsinki.

## Notes

### Competing Interest Statement

The authors have declared no competing interest.

