## Supplementary Data for "Fast prediction of acidic amino acid sidechain conformations for cryo-EM modeling"

**SI fig1. Sidechain dynamics from MD simulations.** The dynamics of sidechain dihedral (C-Cα-Cβ-Cγ) obtained from MD simulations mouse (left) and yeast (right) complex I. Structural values of dihedrals are marked with pink (*Mm*) and cyan (*Yl*) dotted lines.

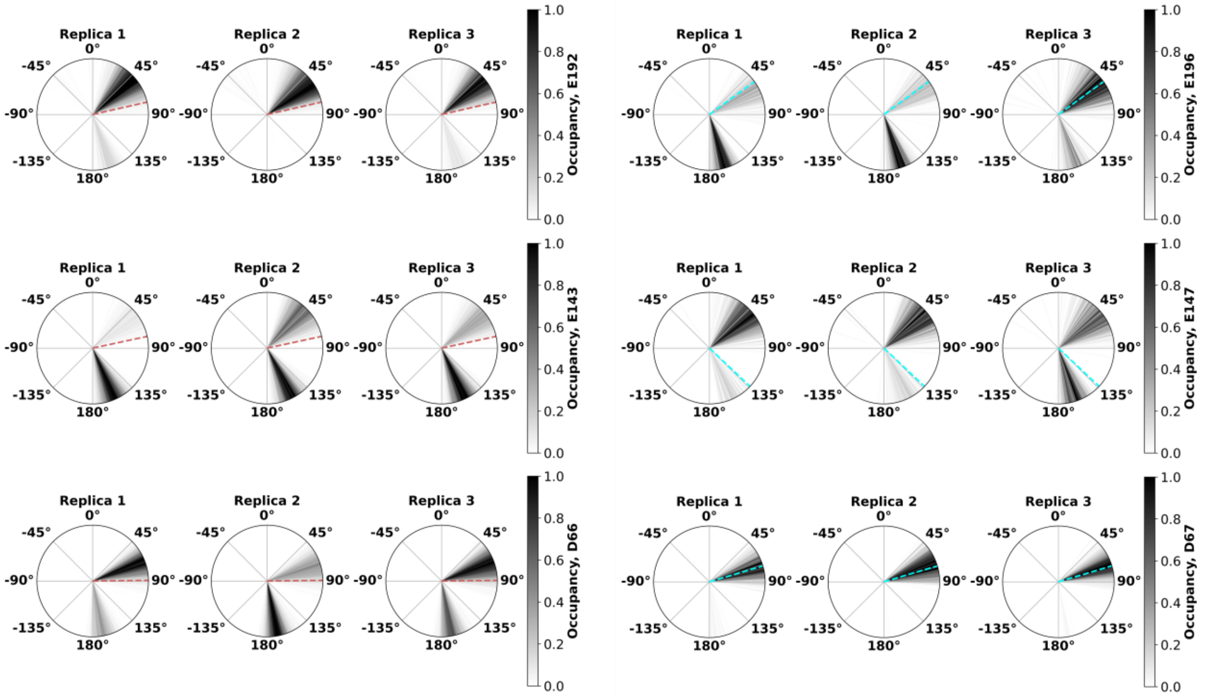

**SI fig2 - Sidechain conformations of acidic amino acid residues of mouse complex I based on Modeler-based modeling and MD simulations.** The pKa vs dihedral angle (C-Cα-Cβ-Cγ) plots (left) and dihedral angle dynamics from MD simulations (right) are shown. pKa of structural conformations are shown as orange dot and the grey dotted line corresponds to pKa = 7. Structural dihedral value is shown as a pink dotted line in radial heat maps.

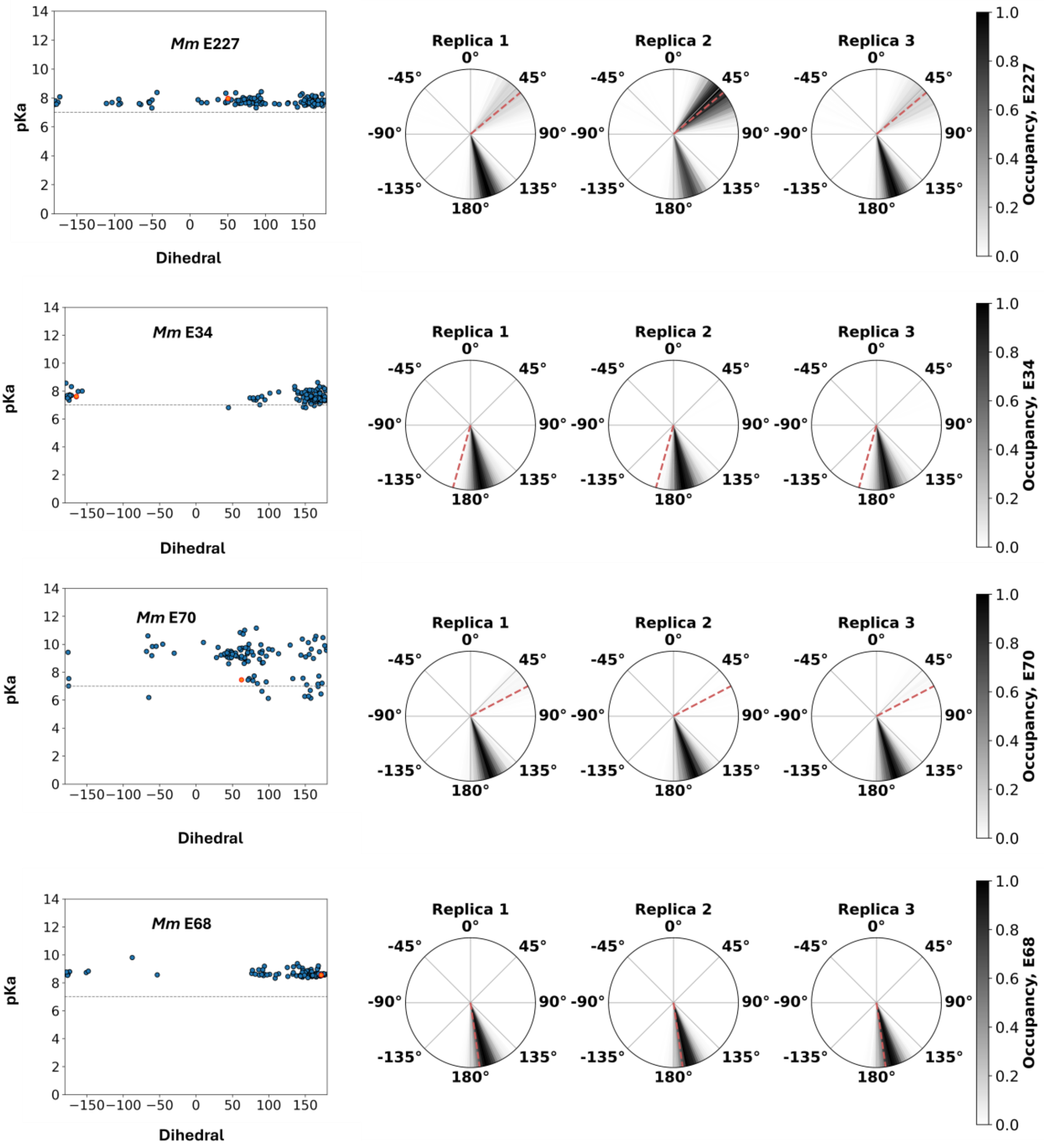

**SI fig3 - Sidechain conformations of acidic amino acid residues from yeast complex I based on Modeler-based modeling and MD simulations.** The pKa vs dihedral angle (C-Cα-Cβ-Cγ) plots (left) and dihedral angle dynamics from MD simulations (right) are shown. pKa of structural conformations are shown as orange dot and the grey dotted line corresponds to pKa = 7. Structural dihedral value is shown as a cyan dotted line in radial heat maps.

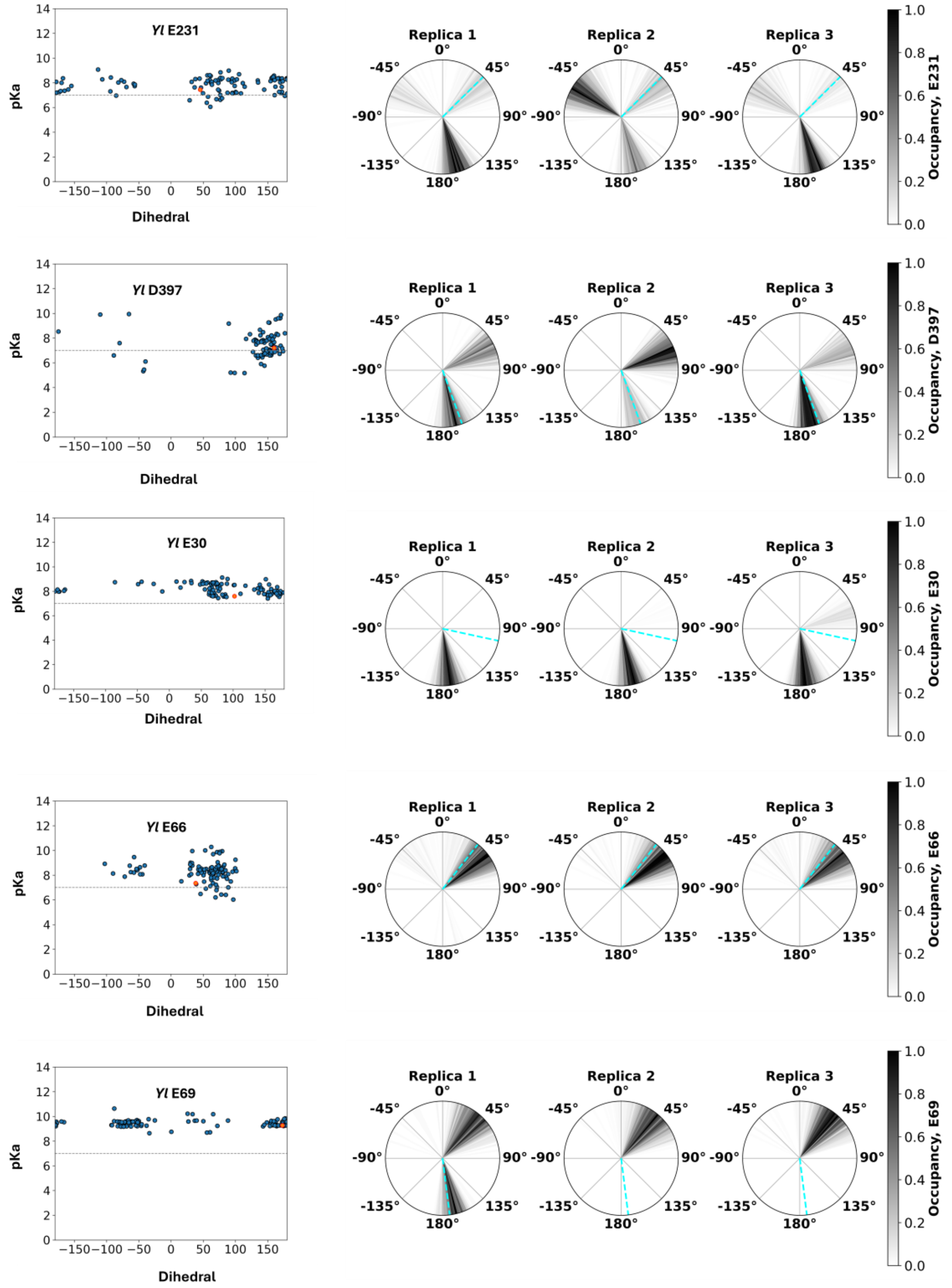
